# Quality assessment of RNA 3D structure models using deep learning and intermediate 2D maps

**DOI:** 10.1101/2025.07.25.666746

**Authors:** Xiaocheng Liu, Wenkai Wang, Zongyang Du, Ziyi Wang, Zhenling Peng, Jianyi Yang

**Affiliations:** MOE Frontiers Science Center for Nonlinear Expectations, Research Center for Mathematics and Interdisciplinary Sciences, Shandong University, Qingdao 266237, China; Chongqing Key Laboratory of Big Data for Bio Intelligence, Chongqing University of Posts and Telecommunications, Chongqing 400065, China; Department of Biomedical Engineering, Chongqing University of Posts and Telecommunications, Chongqing 400065, China

**Author notes:** Equal contribution (Xiaocheng Liu, Wenkai Wang).

## Abstract

Accurate quality assessment is critical for computational prediction and design of RNA three-dimensional (3D) structures. In this work, we introduce RNArank, a novel deep learning-based approach to both local and global quality assessment of predicted RNA 3D structure models. For a given structure model, RNArank extracts a comprehensive set of multi-modal features and processes them with a Y-shaped residual neural network. This network is trained to predict two intermediate 2D maps, including the inter-nucleotide contact map and the distance deviation map. These maps are then used to estimate the local and global accuracy (i.e., lDDT). Extensive benchmark tests indicate that RNArank consistently outperforms traditional methods and other deep learning-based methods. Moreover, RNArank demonstrates promising performance in identifying high-quality structure models for targets from the recent CASP15 and CASP16 experiments. We anticipate that RNArank will serve as a valuable tool for the RNA biology community, improving the reliability of RNA structure modeling and thereby contributing to a deeper understanding of RNA function.

## Introduction

RNA (ribonucleic acid) plays an essential role in gene expression regulation and numerous cellular processes. Its biological function largely depends on its three-dimensional (3D) structure. Unlike proteins, RNA molecules are more structurally flexible and exhibit a higher degree of conformational diversity. The complexity of RNA structure is increased by the weak and long-range interactions that govern RNA folding, making experimental determination of RNA structure more difficult than protein structure^1^.

The inherent challenges of experimental RNA structure determination have long motivated the development of computational RNA structure prediction approaches^2, 3, 4, 5, 6, 7, 8, 9, 10, 11^. In recent years, deep learning-based approaches, in particular, have shown promise in enhancing the accuracy of automated predictions^12, 13, 14, 15, 16, 17, 18, 19, 20^. Nevertheless, unlike the protein field, which has been revolutionized by AlphaFold2^21^, RNA structure prediction currently lacks a comparably high-confidence solution^22^. Community-wide assessments like CASP and RNA-Puzzles consistently show that while automated servers demonstrate competitive results on specific targets, they still generally lag behind human expert groups^23, 24, 25^. Consequently, a hybrid approach combining human intervention with a diverse mix of modeling methods remains essential for modeling the most challenging targets^24^. This context highlights a critical need for RNA structure quality assessment (QA), to rank models generated by both computational and human-led efforts.

Existing QA methods can be broadly classified into two categories: knowledge-based statistical potentials^26, 27, 28^ and deep learning-based approaches^29, 30, 31^. Knowledge-based methods, such as cgRNASP^26^, rsRNASP^27^, and DFIRE-RNA^28^, derive scoring potentials from simulated reference states at either all-atom or coarse-grained levels. However, the accuracy of these methods is often constrained by the incomplete understanding of the principles governing RNA energetic stability, which can lead to the use of inaccurate potentials and reference states that diverge from native conformations.

To address the shortcomings of knowledge-based potentials, deep learning-based methods were introduced, leveraging supervised learning to directly predict the accuracy of RNA structure models^29, 30, 31^. Early deep learning approaches like RNA3DCNN^30^ employed 3D convolutional neural network on a voxelized representation of the 3D structure, while ARES^29^ utilized an Equivariant Graph Neural Network (EGNN). However, the performance of these methods was often hampered by their reliance on limited training models (e.g., from molecular dynamics simulations or Rosetta fragment assembly) and the use of RMSD as a supervision signal. The superposition- and length-dependent nature of RMSD increased the difficulty of learning and limited their generalization. More recently, lociPARSE^31^, inspired by AlphaFold2 ^21^, used a modified Invariant Point Attention (IPA) module to directly predict the lDDT score^32^.

In this work, we present RNArank, a novel deep learning-based approach to both local and global QA. From a multi-modal description of the input structure, RNArank employs a Y-shaped stack of residual neural network (ResNet^33^) blocks to predict two intermediate 2D maps, which are then used to derive the local and global accuracy in terms of lDDT. Benchmark tests demonstrate that RNArank excels at both assessing overall structure quality and selecting high-quality models.

## Results

### Overview of RNArank

The architecture of the RNArank is shown in Fig. 1. For a given RNA structure model, RNArank extracts a comprehensive set of multi-modal features, including 1D features (nucleotide sequence, backbone orientations, Ultrafast Shape Recognition (USR)^34^), 2D features (inter-nucleotide distances, Rosetta two-body energies, clash probabilities), and 3D features (nucleotide-level voxelization). A detailed description of these features is provided in Table S1. These features are processed by a multi-modal encoder to obtain a unified representation (Fig. 1b). Inspired by protein structure assessment methods^35, 36^, rather than predicting a single scalar value, we utilize a Y-shaped residual neural network (Y-ResNet) to predict two intermediate 2D maps, i.e., a contact map and a distance deviation map, which are the core components of the lDDT calculation. These maps are then used to predict the per-residue and global lDDT score (denoted by pLDDT). A detailed description of the model architecture and training procedures can be found in the Methods section. Furthermore, lDDT was originally defined for protein structure, which is not necessarily appropriate for measuring RNA structure quality. To better reflect the structural properties of RNA, we adapted the standard lDDT (denoted as lDDT_raw_) formulation by using larger distance thresholds (see Methods, Eq. (1)). This RNA-adapted lDDT (denoted as lDDT_RNA_) shows a significantly improved correlation with three other well-established metrics (RMSD, TM-score_RNA_, and GDT_TS, see Figs. S1 and S2). Unless otherwise noted, RMSD (calculated via the program RNAalign^37^) and lDDT_RNA_ (Eq. (3, 4)) values are both based on C4’ atoms.

**Figure 1.**
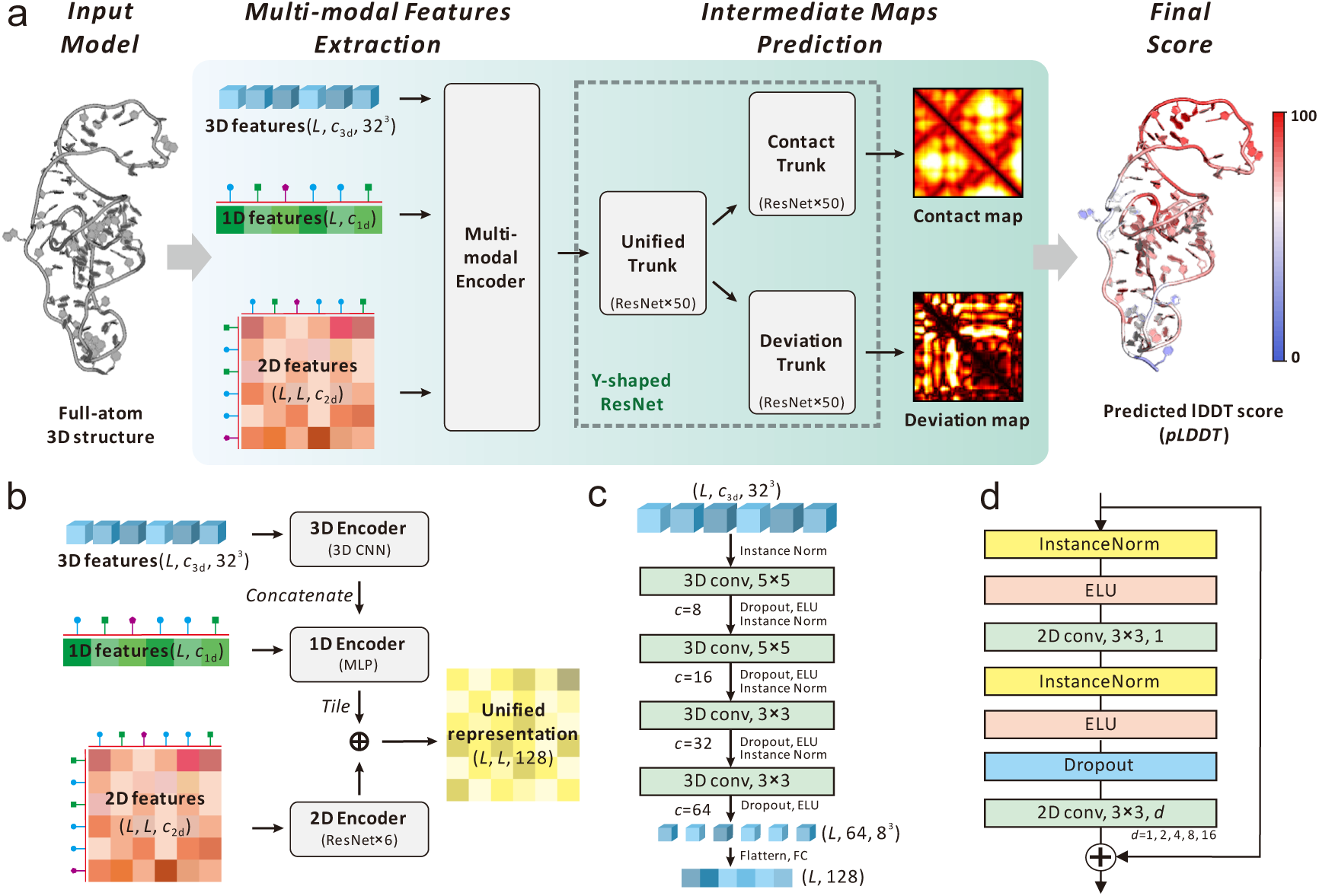
Overall architecture of RNArank. (a) The quality assessment pipeline of RNArank. An input RNA structure model is first processed to extract multi-modal features (1D, 2D, and 3D; with dimensions of *c*_1d_=67, *c*_2d_=58, and *c*_3d_=3, respectively). After encoding and aggregating, these features are fed into a Y-shaped ResNet that predicts two intermediate outputs (a contact map and a deviation map) to derive the final per-nucleotide pLDDT score. The global pLDDT score is calculated by averaging the scores over all nucleotides. (b) The multi-modal encoder. The 3D features are first processed by a 3D CNN block, after which they are flattened and concatenated with the 1D features, then updated by an MLP. In parallel, the 2D features are processed by six ResNet blocks. Finally, the updated 1D representation is tiled to a 2D pairwise map and combined with the processed 2D features via element-wise addition to obtain a unified 2D representation. (c) The 3D CNN block. The 3D features are processed by a stack of 3D convolutional layers (where *c* denotes the number of output channels), flattened, and then mapped to a 1D representation via a fully connected (FC) layer. (d) The ResNet block used in RNArank. We employ a "full pre- activation" architecture^46^ and incorporate dilated convolutions (with a variable rate, *d*) to enhance performance.

### Performance on 24 independent RNA targets

To evaluate the performance of RNArank, we collected a non-redundant dataset of 24 RNAs from PDB (named T24), all released after the date of the training RNAs (January 2022). For each RNA, 35 structure models were generated using various RNA structure prediction methods, including deep learning-based and physics-based approaches^9, 12, 13, 14, 15, 16, 17^ (see Methods and Table S2).

We compare RNArank with five representative QA methods, including three traditional knowledge-based methods (cgRNASP^26^, rsRNASP^27^, DFIRE^28^), and two deep learning-based methods (RNA3DCNN^30^, lociPARSE^31^). The knowledge-based methods provide global QA scores only, while the deep learning-based methods can give both global and local QA scores. Note that lociPARSE is designed to predict lDDT_raw_.

Multiple metrics are used for performance evaluation, including Spearman correlation (*ρ*), accuracy of top-ranked model (lDDT_RNA_ and RMSD), and receiver operating characteristic (ROC). The Spearman correlation (*ρ*_lDDT(Global)_, *ρ*_lDDT(Local)_, and *ρ*_RMSD_) is defined between the predicted QA scores and the true values at both global (molecule; for lDDT_RNA_ and RMSD) and local (nucleotide; only for lDDT_RNA_) levels. These correlations are analyzed at two different levels: on an *overall* basis, by pooling all decoys together; and as a *per-target* basis, by first calculating the correlation for each target and then averaging the results. The accuracy of the top-ranked model (lDDT_RNA_ and RMSD) is defined on the *per-target* basis; while the ROC and the area under the curve (AUC) are calculated on the *overall* basis, to distinguish high-quality models from the rest (see Methods section).

As shown in Table S3, RNArank consistently outperforms existing methods across all evaluation metrics. Notably, despite being trained solely on lDDT_RNA_, RNArank generalizes robustly to RMSD-based evaluations (Fig. 2a). Regarding global molecule-wise assessment, it achieves the highest overall *ρ*_lDDT(Global)_ (0.770 vs. 0.557 for the second-best method) and *ρ*_RMSD_ (0.675 vs. 0.414 for the second-best). This lead is maintained on a *per-target* basis for both lDDT_RNA_ (0.577 vs. 0.551 for the second-best) and RMSD (0.435 vs. 0.272 for the second-best). This strong performance extends to local residue-wise assessment, where RNArank’s overall (0.823) and per-target average (0.644) correlations significantly exceed the other methods (0.581 and 0.499 for the second-best, respectively). These results demonstrate the state-of-the-art performance of RNArank for both global and local QA.

**Figure 2.**
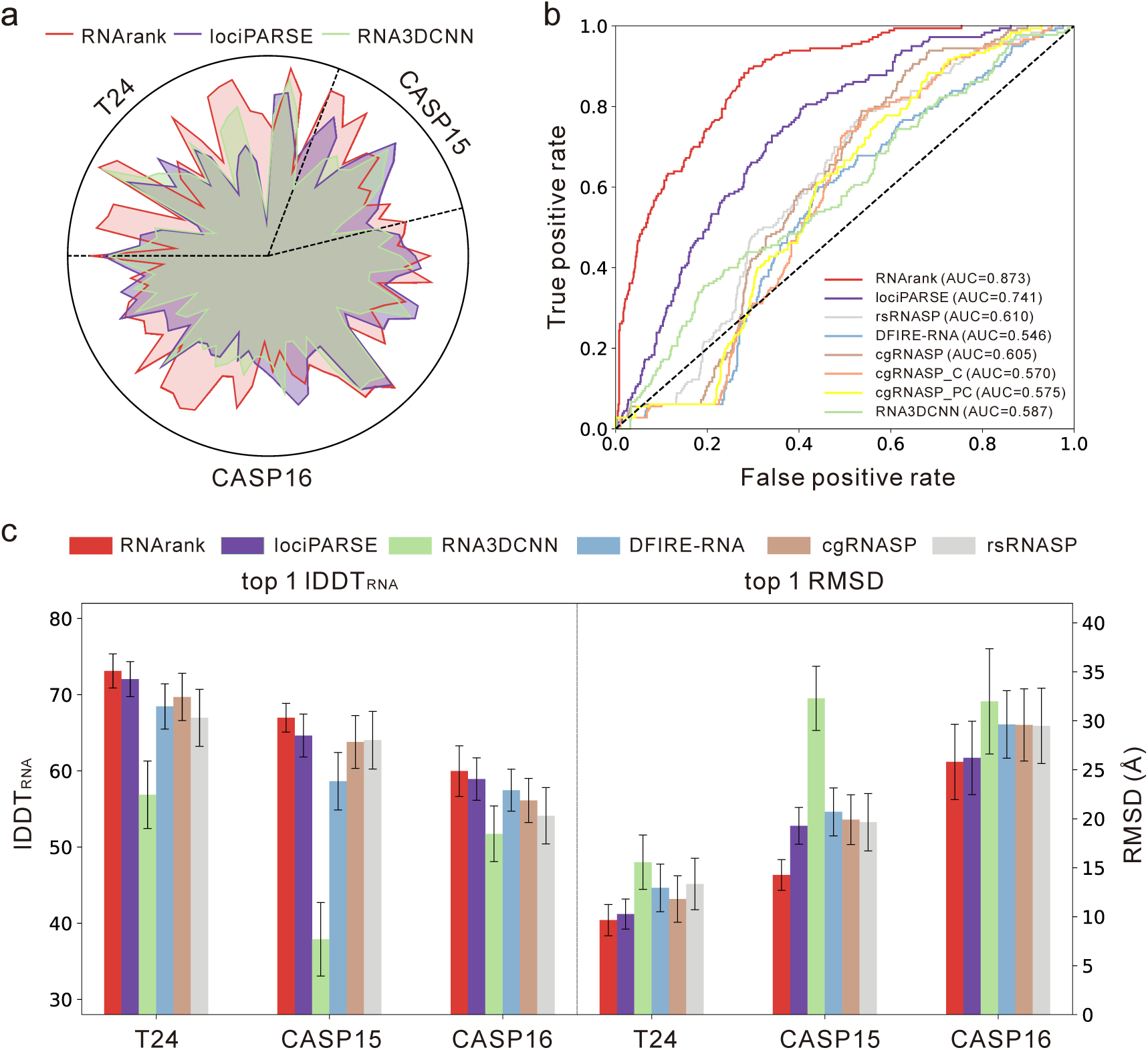
Performance of RNArank and comparison with other methods. **(a)** Comparison of the RMSD Spearman correlation (*ρ*_RMSD_) against lociPARSE (lDDT-based) and RNA3DCNN (RMSD-based) for all tested targets. The radial axis is normalized such that a point further from the center always indicates a better score. (b) ROC curves of predicted scores for structure models from the T24 dataset. The threshold for the positive class is set at a true lDDT_RNA_ score of >75. Note that the scores from knowledge-based methods and RNA3DCNN were inverted before calculating the ROC curves. (c) Comparisons of the top 1 lDDT_RNA_ (left) and RMSD (right) on the three test sets. Error bars represent 20% of the standard deviation.

A crucial aspect of QA is the ability to identify high-quality models. We assess this capability in two ways. First, in the ROC analysis to distinguish high-quality models (lDDT_RNA_ > 75 for positives; Fig. 2b and Table S3), RNArank achieves a leading AUC of 0.873, substantially higher than other methods (0.741 for the second best). Second, we further examine the capacity to identify the best model (i.e., top 1 selection) for each target. As shown in Table S3 and Fig. 2c, RNArank yields the average top 1 lDDT_RNA_ of 73.1 and RMSD of 9.6 Å, better than the second-best method (72.0 and 10.3 Å, respectively). These results demonstrate RNArank’s strengths in model selection. Fig. 3 highlights the reliable quality assessment by RNArank on a representative example (PDB ID: 7TZS; Fig. 3a). On this example, the scores predicted by RNArank correlate strongly (*ρ>*0.9) with the true lDDT_RNA_ values at both the global (Fig. 3b) and local levels (Fig. 3c). This high accuracy is consistently observed across a diverse set of models with varying quality, generated by methods including AlphaFold3, trRosettaRNA, and SimRNA (Figs. 3d-f). In summary, RNArank achieves satisfactory performance in RNA 3D structure QA on the T24 benchmark. This performance marks the advancement in RNA model evaluation made by the RNArank framework.

**Figure 3.**
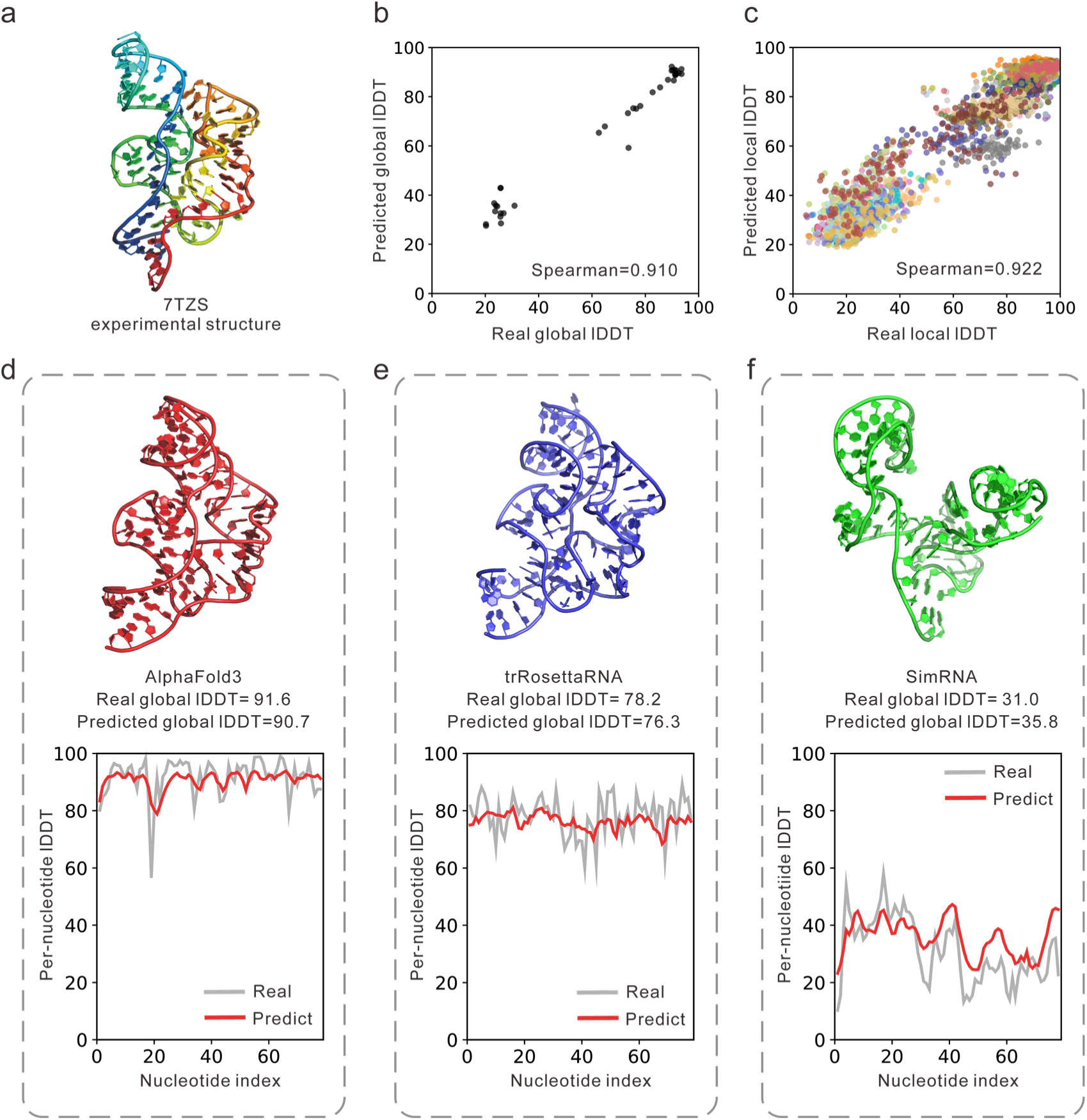
Quality assessment by RNArank for an example target (PDB ID: 7TZS). (a) Experimental structure of 7TZS. (b, c) Correlation between predicted and true lDDT-RNA scores at the (b) global and (c) per-nucleotide (local) levels. Each point in (c) corresponds to a nucleotide, and all nucleotides from the same model are shown with the same color. (d-f) The local lDDT_RNA_ curves for three representative structure models from AlphaFold3, trRosettaRNA, and SimRNA.

### Performance on CASP15 RNAs

The CASP15 in 2022 introduced the RNA structure prediction track, reflecting growing interest in RNA structure modeling. The 12 RNA targets in CASP15 (8 natural, 4 synthetic) presented significantly greater challenges compared to the T24 set, evidenced by a higher and wider RMSD range (Fig. S3). The corresponding 1,622 decoys were generated by a diverse range of structure prediction methods, from new deep learning techniques like trRosettaRNA^14^ (Yang-Server), RhoFold^17^ (AIchemy_RNA), RoseTTAFoldNA^15^ (BAKER), and DeepFoldRNA^12^ (DF_RNA) to traditional approaches such as BRIQ^38^ (AIchemy_RNA2), Vfold^39^ (Chen), RNAComposer^40^ (RNApolis), and SimRNA^9^ (GeneSilico), as well as the interventions from human experts. This methodological diversity ensures a more comprehensive evaluation of RNArank’s performance. Furthermore, these targets were released after the date of training RNAs and are non-identical with our training dataset (see Methods), making them a rigorous benchmark for RNArank’s performance. As shown in Table S4, RNArank consistently outperforms other methods across 8 of the 9 evaluated metrics in the CASP15 test dataset. On the *overall* basis, it achieves Spearman correlation of 0.706 (with global lDDT_RNA_) and 0.686 (with RMSD), significantly outperforming the second-best method (0.511 and 0.200, respectively). RNArank also achieves a substantially higher AUC of 0.915 compared to others (<0.7), indicating a robust capacity for distinguishing high-quality decoys (Fig. S4a). This high performance extended to the local nucleotide-level assessment (local lDDT_RNA_), where RNArank reaches the highest Spearman correlation among all the methods.

On the *per-target* basis, RNArank still outperforms other methods on the RMSD and local lDDT_RNA_ correlation; however, its global lDDT_RNA_ correlation is slightly lower than lociPARSE, which is primarily due to the inaccurate assessment for four targets: R1149, R1156, R1189, and R1190. Among these challenging targets, R1149 and R1156 are known to be highly dynamic and adopt various conformations. The other two targets, R1189 and R1190, are involved in the complicated interaction with more than 4 protein chains, which may distort their assessments as the sole RNA. Analysis of these four targets (Fig. S5) reveals that RNArank tends to give high scores for the models submitted by TS081 (RNApolis group in CASP15, marked by red stars in Fig. S5), which uses RNAComposer, a method designed for RNA-only modeling that performed imperfectly on these four targets. This suggests that RNArank is challenged by the conformational diversity of dynamic RNAs (R1149, R1156), and may also favor unbound conformations over native bound states for targets like R1189 and R1190.

Nevertheless, when focusing on the critical task of identifying the single best decoy (Fig. 2c), RNArank again achieves the highest average top 1 lDDT_RNA_ (67.0) and RMSD (14.3 Å), outperforming the second-best method (64.6 and 19.3 Å, respectively). These findings confirm that despite the challenges posed by complex or dynamic targets, RNArank’s robust scoring mechanism makes it effective at selecting the high-confidence model from a large and diverse pool of models.

### Performance on CASP16 RNAs

The recent CASP16 competition provided a new and richer RNA benchmark set containing 42 targets and almost 6,500 models, allowing for further rigorous testing of RNArank. This set features a wide range of target types (both monomers and multimers) and complexity levels (58-833 nucleotides). With decoys submitted by 48 human and 16 server groups, the CASP16 set displays even greater diversity than previous benchmarks (Fig. S3). As shown in Fig. 2b and Table S5, RNArank continues to demonstrate strong performance across various metrics (7 of 9) on the CASP16 dataset, further highlighting its robustness.

Notably, RNArank excels in identifying an ensemble of high-quality decoys. This is a particularly relevant capability, as blind assessment experiments like CASP and RNA-Puzzles require participants to submit 5∼10 models for each target. This performance is quantified in Fig. S6, which plots the highest lDDT_RNA_ score found within the top-*k* selected decoys. RNArank (red line) maintains a clear lead over its competitors across all values of *k*. For example, RNArank achieves a top-5 lDDT_RNA_ of 62.8 on average, compared to 60.6 of the second-best method. This result suggests the potential of RNArank for application in blind test experiments such as CASP and RNA-Puzzles. In fact, its integration into our automated server, Yang-Server, contributed to our group’s top performance in the automated RNA structure prediction track of the CASP16 blind test. However, we found that RNArank’s overall Spearman correlation for RMSD (*ρ*_RMSD_=0.722) was slightly below that of the energy-based method DFIRE-RNA (0.751), a trend not observed in the previous two test sets. This performance gap stems from two factors: (1) RMSD’s intrinsic strong correlation with sequence length, and (2) the different sensitivities of scoring methods to sequence length. Firstly, RMSD values naturally increase for larger structures. E.g., for CASP16 targets, RMSD shows a clear correlation with sequence length (Fig, S7a; Spearman correlation: 0.811). Similarly, energy-based methods (such as DFIRE-RNA, rsRNASP, and cgRNASP) display strong negative correlations between their scores and sequence length (Figs. S7d-f; Spearman correlations: -0.943, -0.918, -0.931, respectively), which is consistent with the RMSD-length relationship. In contrast, the lDDT score predicted by RNArank is less influenced by sequence length, with a lower Spearman correlation of 0.594 (Fig. S7b). This difference in length dependency influences the *overall* evaluation, which mixes all targets of different lengths together for the correlation calculation. The significantly greater target length variation in CASP16 (357.7 ± 264.8) than the other two sets (T24: 106.5 ± 100.3; CASP15: 208.5 ± 186.5) further magnified this effect, explaining the distinct trend observed for RMSD correlation in the CASP16 set. Nevertheless, when evaluated on the more rigorous *per-target* basis, RNArank maintains its leading performance in terms of *ρ*_RMSD_, though no method, whether trained on lDDT or RMSD, achieves a value >0.4.

### Ablation study

In this section, we conduct an ablation study to evaluate the impact of four key features on RNArank’s performance: ultrafast shape recognition (USR), voxelization, Rosetta two-body energy, and inter-nucleotide distances. For each feature, we removed it from the input feature sets and re- trained an ablated model using the same training dataset and configuration. The performance of these ablated models was benchmarked against the fully-featured RNArank on the T24 dataset, with results summarized in Fig. 4 and Table S6.

**Figure 4.**
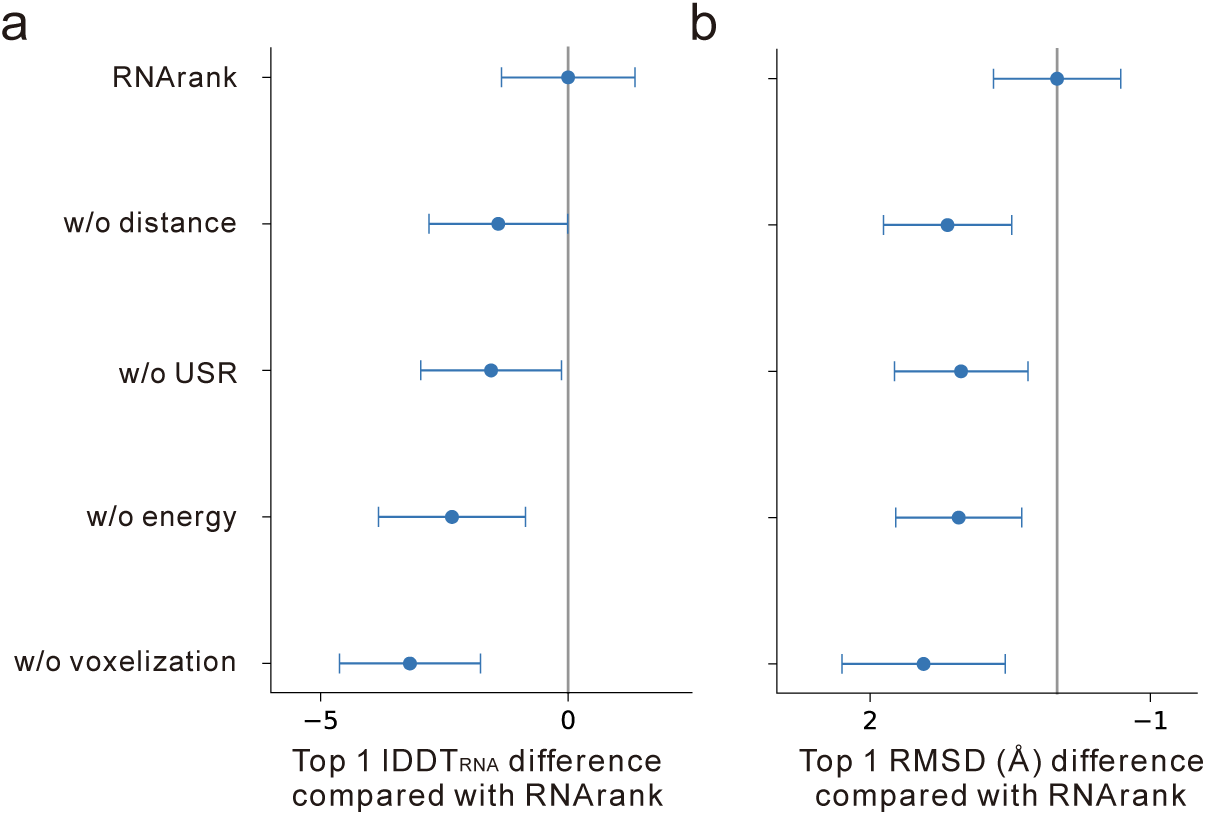
Ablation study on the T24 dataset. Comparison of the per-target average top 1 lDDT_RNA_ (a) and RMSD (b) between the full RNArank model and its ablated variants. Error bars represent 10% of the standard deviations.

As anticipated, the most drastic performance degradation is caused by the removal of the voxelization features, with the average top 1 lDDT_RNA_ dropping by 3.6% (from 73.1 to 70.5) and the average top 1 RMSD increasing by 14.9% (from 9.6 to 11.1 Å). This highlights that while 1D and 2D features provide helpful information, the explicit 3D atomic environment captured by voxelization is indispensable for fine-grained quality assessment. Interestingly, we observed that removing USR or voxelization unexpectedly improved several *overall* (i.e., dataset-wide) metrics. We attribute this to the high degree of heterogeneity in topology and local environments across different RNA targets. When all decoys are pooled for a single metric, these highly target-specific features may hinder the model’s ability to find a universal quality signal, thus acting as a source of noise. Nevertheless, when performance is evaluated on a more meaningful *per-target* basis, removing these features consistently degrades performance across both score correlations and top 1 accuracy. This ultimately confirms the critical importance of both USR and voxelization features for robust quality assessment.

Finally, ablating the inter-nucleotide distance and Rosetta energy features also led to notable performance decreases, particularly in top 1 model selection (top 1 lDDT_RNA_ scores dropped from 73.1 to 72.0 and 71.6, respectively). This demonstrates that these 2D features, which encode crucial geometric constraints and physicochemical interactions, are helpful for accurately identifying the high-quality models. Taken together, these results validate our multi-modal feature strategy and underscore its importance for robust structure quality assessment.

## Discussion

RNA 3D structure prediction has drawn increased interest in recent years, and the reliable quality assessment plays a critical role in improving structure prediction accuracy. To achieve this, we have developed an automated approach for RNA 3D structure quality assessment: RNArank, which utilizes comprehensive multi-modal features and a Y-shaped ResNet to estimate the lDDT of a predicted structure model. We rigorously benchmarked RNArank on three datasets, including 24 independent RNAs and two sets from CASP15 and CASP16. The results show that RNArank consistently achieves promising performance in selecting decoys and outperforms other approaches, including both the traditional energy-based methods and the recent deep learning-based methods.

However, benchmarking on CASP datasets demonstrates that it remains challenging to assess models for RNAs that form complicated multimers and/or interact with proteins. Even with promising methods for assessing docking results like DRPScore^41^, fully accounting for the conformational changes induced by protein binding remains a significant challenge that may hinder the accurate evaluation of complexes from *de novo* predictors like AlphaFold3. Given RNA’s natural tendency to rely on intermolecular interactions for structural stability, the next-generation RNA QA methods must evolve to effectively incorporate information from binding partners and consider the dynamics of complex formation.

## Methods

### lDDT_RNA_ definition

The definition of lDDT in this work (lDDT_RNA_) introduces two critical modifications to the original lDDT (lDDT_raw_) that was designed for proteins. First, we adjusted the set of deviation cutoffs. While lDDT_raw_ uses thresholds of {0.5 Å, 1.0 Å, 2.0 Å, 4.0 Å}, lDDT_RNA_ employs a shifted set of {1.0 Å, 2.0 Å, 4.0 Å, 6.0 Å}. This change accounts for the larger distance scale inherent to RNA structures.

Second, we extended the inclusion radius for contact definition from the standard 15 Å (used in lDDT_raw_) to 30 Å. This expansion is crucial to accommodate the larger size and more extended geometry of nucleotides; for instance, the C4’-C4’ distance between base-paired nucleotides can exceed the standard 15 Å cutoff. Specifically, the lDDT_RNA_ score is formulated as:

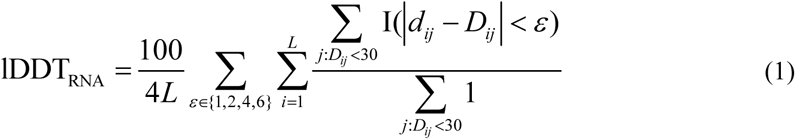

where *L* is the total number of nucleotides; *d_ij_* and *D_ij_* are the C4’ distance between *i*-th and *j*-th nucleotides in the predicted and experimental structures, respectively; I() is the indicator function.

The effectiveness of these modifications was confirmed by comparing with other established metrics. We calculated the Spearman correlation coefficients between both lDDT versions and three established quality metrics, RMSD, TM-score_RNA_, and GDT_TS, across the CASP15 and CASP16 datasets (Figs. S1 and S2). The analysis reveals a clear and consistent improvement for lDDT_RNA_ in nearly all cases. Overall, lDDT_RNA_ provides a more robust and reliable assessment of RNA model quality than the standard lDDT formulation.

### RNArank algorithm

As shown in Fig. 1a, the RNArank pipeline involves three key steps: (1) preparation of the multi- modal input features, (2) prediction of contact and deviation maps using a Y-shaped residual neural network, and (3) calculation of the quality assessment score pLDDT. An overview of the network architecture and feature types is presented below, and a more detailed description of the features is available in Table S1.

#### Step 1. Preparation of input features

The input of RNArank is the full-atom structure of the model to be assessed. Three sets of features are calculated from this input as follows:

**1D features.** This set includes the RNA sequence, local backbone geometry, and global shape information from ultrafast shape recognition (USR). The sequence is represented by one-hot encoding of the four nucleotide types, while local geometry is defined by the seven backbone torsion angles. To capture global topology in per-nucleotide view, we employ the USR method, which has demonstrated its effectiveness in protein QA^34, 36^. In RNArank, USR features are calculated using the C4’ atom as the representative point for each nucleotide, describing the overall structure via per-nucleotide topological moments.

**2D features.** This set primarily describes the inter-nucleotide relationships, comprising Euclidean distances, Rosetta two-body centroid energy terms, and the steric clashes (Eq. (2)). Inter- nucleotide distances are calculated between the C4’, P, and N atoms, encoded using Gaussian radial basis function. The Rosetta energy features consist of various two-body terms, including fa_atr, fa_rep, lk_nonpolar, rna_torsion, fa_stack, stack_elec, geom_sol_fast, hbond_sc, and fa_elec_rna_phos_phos. These energy terms are normalized before being fed into the neural network. The steric clashes are quantified by calculating the frequency of non-hydrogen atom pairs whose distance falls below a threshold determined by their van der Waals radii. The specific formula for this clash metric is as follows:

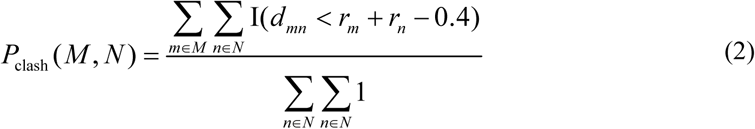

where *d_mn_* is the distance between atoms *m* and *n*, which belong to nucleotides *M* and *N*, respectively. *r_i_* is the van der Waals radius of atom *i*.

**3D features.** This set consists of the voxelized representation of each nucleotide’s local atomic environment, using a method similar to RNA3DCNN^30^. To ensure rotational and translational invariance, we first establish a local coordinate system for each nucleotide based on its P, C4’, and key base atoms (N1 for pyrimidines; N9 for purines). The surrounding environment is then mapped onto a 3D grid aligned with this local frame. Each grid box represents a voxel with three channels, which store the summed atomic occupation, mass, and charge, respectively. This process transforms each nucleotide’s surrounding environment into a 3D representation that is invariant to global translations and rotations.

The above three sets of input features are encoded through a multi-modal encoder consisting of separate convolutional layers corresponding to their respective dimensionalities (Fig. 1b). Specifically, the 3D voxelization features are passed through 3D convolutional layers, flattened, and then concatenated with the 1D features; the resulting tensor is subsequently encoded by 1D convolutions, tiled to match the 2D feature dimensions, and added to the 2D features. Finally, this combined tensor is fed into a 2D ResNet block to generate a unified 2D representation.

#### Step 2. Prediction of 2D contact and deviation maps

In the second step, the unified representation obtained in step 1 is fed into a Y-shaped ResNet module to predict the contact and deviation maps. The contact map is defined as the inter-nucleotide C4’ distance below 30 Å. The deviation map is defined as the one-hot encoding of the distance error falling into five bins corresponding to the deviation cutoffs: <1 Å, 1∼2 Å, 2∼4 Å, 4∼6 Å, and >6 Å. Specifically, this module consists of three stages:

1. **Unified representation updating.** The initial unified representation is updated using a stack of 50 ResNet blocks. Notably, dilated convolutions (Fig. 1c) are incorporated to improve the efficiency of capturing long-range dependencies while maintaining computational efficiency.
2. **Task-specific representation learning.** The output from the first stage is then channeled into two parallel branches: one with 50 ResNet blocks to learn the contact representation, and another with 50 ResNet blocks for the deviation representation.
3. **Output layer.** Finally, each representation is passed through a linear layer followed by either a sigmoid (for contact prediction) or a softmax (for deviation prediction) activation to generate the predicted probability maps.

#### Step 3. Calculation of the predicted lDDT (pLDDT) score

In the final step, the predicted 2D contact and deviation maps are used to calculate the pLDDT score. The calculation proceeds as follows. For any pair of nucleotides (*i*, *j*), the two predicted maps can derive: (1) contact probability,, which is the model’s confidence that the C4’-C4’ distance between these nucleotides are within 30 Å; (2) deviation probabilities, *P^ij^*, representing the probability that the deviation in the C4’-C4’ distance is less than a given threshold *d*∈{1.0 Å, 2.0 Å, 4.0 Å, 6.0 Å}. Subsequently, the local pLDDT score for the *i*-th nucleotide is calculated following the lDDT_RNA_ definition (eq. (1)). The global pLDDT score is then computed by averaging all the local scores. The specific formulas are as follows:

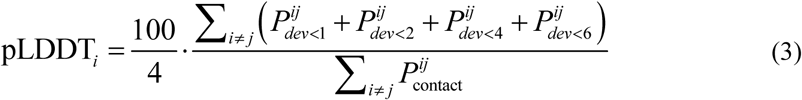

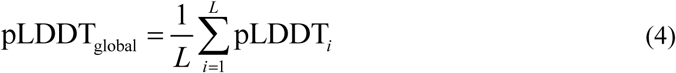

where *L* is the sequence length.

### Construction of datasets

**Benchmark datasets.** Our evaluation is based on three distinct benchmark datasets. The first set (T24) contains 24 RNA-only entries from PDB (released after January 2022) for which we generated a comprehensive model library using a variety of RNA structure prediction methods, including AlphaFold3^16^, DeepFoldRNA^12^, DRFold^13^, RhoFold^17^, RoseTTAFold2NA^15^, SimRNA^9^, and trRosettaRNA^14^. The number of structure models generated by each method is listed in Table S2. Two additional test sets were derived from the CASP15 and CASP16 competitions, containing 12 and 42 targets, respectively. For these two sets, we assessed the original models submitted by the official participants, providing a realistic and challenging benchmark.

**Training sets.** To train our models, we initially collected RNAs from the trRosettaRNA training dataset, which included experimental structures resolved before January 2022. This dataset was then filtered to remove RNAs with non-standard nucleotides, chain discontinuities, or formatting errors in their coordinate files. We also excluded sequences longer than 200 nucleotides. To ensure a strict separation from our test sets, we used cd-hit-est^42^ to remove any sequences identical to those in our test sets. This filtering process resulted in a final set of 1,635 RNAs.

For each of these 1,635 RNAs, we generated a diverse set of structure models, totaling approximately 200,000 models. These models were generated using a combination of various strategies: (1) deep learning-based prediction, (2) physics-based folding, (3) structure perturbation, (4) molecular dynamics simulations, and (5) fragment assembly-based modeling. The details of each approach are outlined below.

1. **Deep learning prediction.** AlphaFold3, DeepFoldRNA, DRFold, RoseTTAFold2NA, RhoFold, and trRosettaRNA were installed and run locally on our computing cluster.
2. **Physics-based folding.** We also employed SimRNA, a physics-based RNA folding method that simulates the folding process based on the physical interactions between nucleotides, providing a complementary approach to deep learning-based methods.
3. **Structure perturbation.** For each experimental structure, we applied random perturbations to the backbone angles of each nucleotide using a standard normal distribution. These perturbations allow for the generation of diverse decoys while maintaining structural plausibility, providing a more comprehensive dataset for training the model.
4. **Molecular dynamics simulations.** We performed molecular dynamics (MD) simulations using GROMACS^43^ to model RNA structural dynamics. For each RNA experimental structure, we solvated the system with water and neutralized it by adding ions. The CHARMM36-jul2022 force field was applied, and energy minimization was followed by NVT and NPT equilibration to relax the system. A production MD simulation was then conducted to generate trajectories, with snapshots extracted at regular intervals for analysis.
5. **Rosetta fragment assembly-based modeling.** We employed Rosetta to generate decoys using various templating strategies, including native fragments and templates identified through searches. Additionally, we also used secondary structures predicted by RNAfold^44, 45^ to generate structure models.

To ensure structural diversity and reduce redundancy within the generated structure models, we performed a clustering step. Specifically, for each RNA, we applied k-means clustering to group the models according to their pairwise lDDT_RNA_ matrix, and selected representative models from each cluster to form the final training set. On average, we generated approximately 120 structure models for each RNA, with the explicit number depending on the success of the MD simulations.

### Loss function and training procedure

The ground-truth labels for training include contact maps, deviation maps, and per-nucleotide lDDT scores of each structure model. Correspondingly, the total loss function guiding the training of RNArank is a weighted sum of three components: a contact loss, a deviation loss, and an lDDT loss.

The overall objective is to minimize:

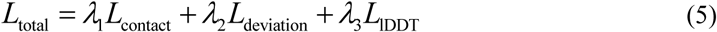

where the weighting factors *λ*_1_, *λ*_2_, and *λ*_3_ were empirically set to 1.0, 1.0, and 20.0, respectively. The individual loss components are defined as follows:

1. ***L*_contact_**: A binary cross-entropy loss of the contact map prediction, which classifies nucleotide pairs as being in contact (< 30 Å) or not.
2. ***L*_deviation_**: A categorical cross-entropy loss of the deviation map prediction, which is framed as a multi-class classification problem over discrete distance error bins. This loss is calculated only for nucleotide pairs that are in contact (i.e., distance < 30 Å) in the experimental structure, thereby focusing the model on learning distance errors for interacting pairs.
3. ***L*_lDDT_**: A mean squared error (MSE) loss of the final per-nucleotide pLDDT score, encouraging the network to accurately regress the true lDDT value.

RNArank was trained using the Adam optimizer with an initial learning rate of 5e-4, an exponential decay rate of 0.99 per epoch, and a dropout rate of 0.2 applied to the ResNet blocks. Each training epoch involved iterating over all RNAs in the training set, with one structure model randomly sampled for each RNA. The model checkpoint with the lowest validation loss (10% of the training samples were retained for validation) was saved for final evaluation. To enhance prediction robustness, we trained two neural network models. The final RNArank score is the average of the pLDDT outputs from these two models, providing a more stable and reliable prediction.

### Selected baseline methods

We benchmark RNArank against a comprehensive suite of existing methods, including traditional knowledge-based statistical potentials (cgRNASP^26^, rsRNASP^27^, and DFIRE-RNA^28^) and deep learning-based methods (RNA3DCNN^30^ and lociPARSE^31^). rsRNASP and cgRNASP are both residue-separation-based statistical potentials. They differ primarily in their granularity; rsRNASP considers all atoms, whereas cgRNASP utilizes three coarse-grained (CG) representations to increase efficiency. In this work, we evaluated all three versions of cgRNASP: the default 3-bead cgRNASP (representing each nucleotide with P, C4’, and N1/N9 atoms for pyrimidine/purine), the 2-bead cgRNASP-PC (P and C4’ atoms), and the 1-bead cgRNASP-C (C4’ atom only). DFIRE-RNA is another all-atom potential, using a distance-scaled finite ideal-gas reference state to derive its energy function.

Among the deep learning-based methods, RNA3DCNN uses a 3D convolutional neural network to predict an RMSD-based “unfitness” score for each nucleotide by treating its local environment as a 3D image. More recently, lociPARSE was developed to estimate the lDDT_raw_ score using the invariant point attention (IPA) architecture, which is inherently invariant to global rotations and translations.

For a fair and rigorous comparison, all baseline methods were installed and run locally on our computing cluster under their default configurations. The sole input for each method was the PDB file of a predicted structure model. For RNA3DCNN, we employed the RNA3DCNN_MD model, which was trained on structure models from molecular dynamics (MD) simulations.

### Performance assessment

The performance of all methods was assessed across multiple criteria to provide a comprehensive evaluation, including correlation with true quality scores, the ability to distinguish high-quality models, and top 1 model selection. The specific metrics are defined as follows:

**Global correlation**. We assess global molecule-wise performance by computing the Spearman’s rank correlation coefficient (*ρ*) between predicted scores and true quality metrics. To provide a comprehensive view, this analysis is conducted on two bases: **(1) *overall* basis**, where the correlation is computed across the pooled set of all structure models from different targets, and **(2) *per-target* basis**, where the correlation is first calculated for each target’s structure models and then averaged across all targets. For both analyses, we evaluate the correlation against two distinct ground-truth metrics: global lDDT_RNA_ and RMSD. We choose Spearman ranking correlation due to its robustness against non-linear relationships and non-normal data distributions.

**Local correlation.** In this case, the correlation is computed between the predicted per-nucleotide scores and the true local lDDT_RNA_ values. Similarly, local-level performance is evaluated on both the *overall* and *per-target* bases.

**Ability to distinguish high-quality models.** This ability is quantified by the AUC calculated on the *overall* basis, with models having a true lDDT_RNA_>75 considered positive samples.

**Top 1 model selection.** To evaluate a QA method’s ability to identify the best model, we report the per-target average accuracy of the top-ranked model in terms of both lDDT_RNA_ and RMSD. A higher average top 1 lDDT_RNA_ and a lower average top 1 RMSD thus indicate better performance.

## Availability

The RNArank web server is available at: https://yanglab.qd.sdu.edu.cn/RNArank/. The source codes are available at: https://github.com/YangLab-SDU/RNArank/.

## Acknowledgements

This work is supported by the following funding sources: National Natural Science Foundation of China (NSFC T2225007, T2222012, 32430063, 62402075), Postdoctoral Fellowship Program and China Postdoctoral Science Foundation (BX20240212), the Science and Technology Research Program of Chongqing Municipal Education Commission (KJQN202300639), and Fundamental Research Funds for the Central Universities.

## Reference

1. Kwon D. RNA function follows form - why is it so hard to predict? Nature 639, 1106–1108 (2025).

2. Madison JT, Everett GA, Kung H. Nucleotide sequence of a yeast tyrosine transfer RNA. Science 153, 531–534 (1966).

3. Levitt M. Detailed molecular model for transfer ribonucleic acid. Nature 224, 759–763 (1969).

4. Sharma S, Ding F, Dokholyan NV. iFoldRNA: three-dimensional RNA structure prediction and folding. Bioinformatics 24, 1951–1952 (2008).

5. Parisien M, Major F. The MC-Fold and MC-Sym pipeline infers RNA structure from sequence data. Nature 452, 51–55 (2008).

6. Rother M, Rother K, Puton T, Bujnicki JM. ModeRNA: a tool for comparative modeling of RNA 3D structure. Nucleic Acids Res 39, 4007–4022 (2011).

7. Zhao Y, Huang Y, Gong Z, Wang Y, Man J, Xiao Y. Automated and fast building of three-dimensional RNA structures. Sci Rep 2, 734 (2012).

8. Popenda M, et al. Automated 3D structure composition for large RNAs. Nucleic Acids Res 40, e112 (2012).

9. Boniecki MJ, et al. SimRNA: a coarse-grained method for RNA folding simulations and 3D structure prediction. Nucleic Acids Res 44, e63 (2016).

10. Watkins AM, Rangan R, Das R. FARFAR2: Improved De Novo Rosetta Prediction of Complex Global RNA Folds. Structure (London, England : 1993) 28, 963–976.e966 (2020).

11. Li J, Zhang S, Zhang D, Chen SJ. Vfold-Pipeline: a web server for RNA 3D structure prediction from sequences. Bioinformatics 38, 4042–4043 (2022).

12. Pearce R, Omenn GS, Zhang Y. De Novo RNA Tertiary Structure Prediction at Atomic Resolution Using Geometric Potentials from Deep Learning. bioRxiv, 2022.2005.2015.491755 (2022).

13. Li Y, Zhang C, Feng C, Pearce R, Lydia Freddolino P, Zhang Y. Integrating end-to-end learning with deep geometrical potentials for ab initio RNA structure prediction. Nat Commun 14, 5745 (2023).

14. Wang W, et al. trRosettaRNA: automated prediction of RNA 3D structure with transformer network. Nat Commun 14, 7266 (2023).

15. Baek M, McHugh R, Anishchenko I, Jiang H, Baker D, DiMaio F. Accurate prediction of protein-nucleic acid complexes using RoseTTAFoldNA. Nat Methods 21, 117–121 (2024).

16. Abramson J, et al. Accurate structure prediction of biomolecular interactions with AlphaFold 3. Nature 630, 493–500 (2024).

17. Shen T, et al. Accurate RNA 3D structure prediction using a language model-based deep learning approach. Nat Methods 21, 2287–2298 (2024).

18. Wang W, Peng Z, Yang J. Predicting RNA 3D structure and conformers using a pre-trained secondary structure model and structure-aware attention. bioRxiv, 2025.2004.2009.647915 (2025).

19. Kagaya Y, et al. NuFold: end-to-end approach for RNA tertiary structure prediction with flexible nucleobase center representation. Nat Commun 16, 881 (2025).

20. Li Y, Feng C, Zhang X, Zhang Y. Ab initio RNA structure prediction with composite language model and denoised end-to-end learning. bioRxiv, 2025.2003.2005.641632 (2025).

21. Jumper J, et al. Highly accurate protein structure prediction with AlphaFold. Nature 596, 583–589 (2021).

22. Schneider B, Sweeney BA, Bateman A, Cerny J, Zok T, Szachniuk M. When will RNA get its AlphaFold moment? Nucleic Acids Res 51, 9522–9532 (2023).

23. Das R, et al. Assessment of three-dimensional RNA structure prediction in CASP15. Proteins 91, 1747–1770 (2023).

24. Kretsch RC, et al. Assessment of nucleic acid structure prediction in CASP16. bioRxiv, 2025.2005.2006.652459 (2025).

25. Bu F, et al. RNA-Puzzles Round V: blind predictions of 23 RNA structures. Nat Methods 22, 399–411 (2025).

26. Tan YL, Wang X, Yu S, Zhang B, Tan ZJ. cgRNASP: coarse-grained statistical potentials with residue separation for RNA structure evaluation. *NAR genomics and bioinformatics* 5, lqad016 (2023).

27. Tan YL, Wang X, Shi YZ, Zhang W, Tan ZJ. rsRNASP: A residue-separation-based statistical potential for RNA 3D structure evaluation. Biophys J 121, 142–156 (2022).

28. Zhang T, Hu G, Yang Y, Wang J, Zhou Y. All-Atom Knowledge-Based Potential for RNA Structure Discrimination Based on the Distance-Scaled Finite Ideal-Gas Reference State. Journal of computational biology : a journal of computational molecular cell biology 27, 856–867 (2020).

29. Townshend RJL, et al. Geometric deep learning of RNA structure. Science 373, 1047–1051 (2021).

30. Li J, et al. RNA3DCNN: Local and global quality assessments of RNA 3D structures using 3D deep convolutional neural networks. PLOS Computational Biology 14, e1006514 (2018).

31. Tarafder S, Bhattacharya D. lociPARSE: A Locality-aware Invariant Point Attention Model for Scoring RNA 3D Structures. Journal of chemical information and modeling 64, 8655–8664 (2024).

32. Mariani V, Biasini M, Barbato A, Schwede T. lDDT: a local superposition-free score for comparing protein structures and models using distance difference tests. Bioinformatics 29, 2722–2728 (2013).

33. He K, Zhang X, Ren S, Sun J. Deep Residual Learning for Image Recognition. In: 2016 *IEEE Conference on Computer Vision and Pattern Recognition (CVPR)*) (2016).

34. Ballester PJ, Finn PW, Richards WG. Ultrafast shape recognition: Evaluating a new ligand-based virtual screening technology. Journal of Molecular Graphics and Modelling 27, 836–845 (2009).

35. Hiranuma N, Park H, Baek M, Anishchenko I, Dauparas J, Baker D. Improved protein structure refinement guided by deep learning based accuracy estimation. Nat Commun 12, 1340 (2021).

36. Guo SS, Liu J, Zhou XG, Zhang GJ. DeepUMQA: ultrafast shape recognition-based protein model quality assessment using deep learning. Bioinformatics 38, 1895–1903 (2022).

37. Gong S, Zhang C, Zhang Y. RNA-align: quick and accurate alignment of RNA 3D structures based on size-independent TM-scoreRNA. Bioinformatics 35, 4459–4461 (2019).

38. Xiong P, Wu R, Zhan J, Zhou Y. Pairing a high-resolution statistical potential with a nucleobase-centric sampling algorithm for improving RNA model refinement. Nature Communications 12, 2777 (2021).

39. Li J, Zhang S, Zhang D, Chen S-J. Vfold-Pipeline: a web server for RNA 3D structure prediction from sequences. Bioinformatics 38, 4042–4043 (2022).

40. Antczak M, et al. New functionality of RNAComposer: an application to shape the axis of miR160 precursor structure. Acta biochimica Polonica 63, 737–744 (2016).

41. Zeng C, Jian Y, Vosoughi S, Zeng C, Zhao Y. Evaluating native-like structures of RNA-protein complexes through the deep learning method. Nat Commun 14, 1060 (2023).

42. Fu L, Niu B, Zhu Z, Wu S, Li W. CD-HIT: accelerated for clustering the next-generation sequencing data. Bioinformatics 28, 3150–3152 (2012).

43. Páll S, et al. Heterogeneous parallelization and acceleration of molecular dynamics simulations in GROMACS. The Journal of chemical physics 153, 134110 (2020).

44. Hofacker IL. Vienna RNA secondary structure server. Nucleic Acids Res 31, 3429–3431 (2003).

45. Lorenz R, et al. ViennaRNA Package 2.0. Algorithms for molecular biology : AMB 6, 26 (2011).

46. He K, Zhang X, Ren S, Sun J. Identity Mappings in Deep Residual Networks. *arXiv e-prints*, arXiv:1603.05027 (2016).

