## Supplementary Material for "Quality assessment of RNA 3D structure models using deep learning and intermediate 2D maps"

### Supplementary Information

#### Catalogue

#### Table S1. Extracted 1D, 2D and 3D features.

Table S1. Extracted 1D, 2D and 3D features. *L* is the sequence length.

| Type | Name and shape | Description |
|---|---|---|
| 1D | Nucleotide sequence<br>(*L*, 5) | One-hot encoded base types for each nucleotide,<br>and encoded relative position information in the sequence. |
| | backbone orientations<br>(*L*, 14) | Seven backbone torsion angles: α, β, γ, δ, ε, ξ and χ;<br>Represented using sine and cosine. |
| | Ultrafast Shape Recognition (USR)[1, 2]<br>(*L*, 48) | For each C4' atom, three sets of distances (from the current C4', from the C4' furthest from the current, and from the C4' furthest from the second reference) to all other C4' atoms are calculated. Each of these three distance sets is then encoded using Gaussian radial basis functions. |
| 2D | inter-nucleotide distance<br>(*L*, *L*, 48) | Three distance maps for each pair of nucleotides:<br>P to P, C4' to C4', and glycosidic N to glycosidic N distance maps;<br>Distances are encoded by Gaussian radial basis functions, respectively. |
| | Rosetta energies<br>(*L*, *L*, 9) | Two-body terms in Rosetta:<br>fa_atr, fa_rep, lk_nonpolar, rna_torsion, fa_stack, stack_elec, geom_sol_fast, hbond_sc, and fa_elec_rna_phos_phos |
| | Steric clashes<br>(*L*, *L*, 1) | The frequency of pairwise non-hydrogen atomics clash:<br>$P_{\text{clash}}(M,N)=\frac{\sum_{m\in M}\sum_{n\in N} \mathrm{I}(d_{mn}<r_m+r_n-0.4)}{\sum_{n\in N}\sum_{n\in N} 1}$<br>The atomic radii for carbon (C), nitrogen (N), oxygen (O), and phosphorus (P) atoms are set as 1.7, 1.55, 1.52, and 1.8 Å, respectively |
| 3D | nucleotide-level voxelization<br>(*L*, 3, 32, 32, 32) | Each nucleotide is represented by a 3-channel (atomic occupation, mass, charge) voxelization of its atoms into a 32 × 32 × 32 Å grid, resulting in a (3, 32, 32, 32) feature map per nucleotide.[3] |

#### Table S2. Number of decoys per RNA target in T24 benchmark sets.

Table S2. Number of decoys per RNA target in T24 benchmark sets.

| Method | Number of decoys per target |
|---|---|
| AlphaFold3[4] | 5 |
| DeepFoldRNA[5] | 12 |
| DRFold[6] | 8 |
| RhoFold[7] | 2 |
| RoseTTAFold2NA[8] | 2 |
| trRosettaRNA[9] | 5 |
| SimRNA[10] | 1 |
| Total | 35 |

#### Table S3. Performance on the test dataset T24.

Values in **bold** indicate the best performance, while those with underlines represent the second-best. AUC was calculated on the overall data with an $\mathrm{lDDT_{RNA}}$ cutoff of 75 to differentiate between 'good' and 'bad' decoys. $\rho_{\mathrm{lDDT(Global)}}$, $\rho_{\mathrm{lDDT(Local)}}$ and $\rho_{\mathrm{RMSD}}$ represent the Spearman correlation between the assessment scores and the true global $\mathrm{lDDT_{RNA}}$, local $\mathrm{lDDT_{RNA}}$, and RMSD, respectively. All correlation values are reported as absolute values.

| Method | Overall | | | | Per-target average | | | | |
|---|---|---|---|---|---|---|---|---|---|
| | $\rho_{\mathrm{lDDT(Global)}}$ | $\rho_{\mathrm{lDDT(Local)}}$ | $\rho_{\mathrm{RMSD}}$ | AUC | $\rho_{\mathrm{lDDT(Global)}}$ | $\rho_{\mathrm{lDDT(Local)}}$ | $\rho_{\mathrm{RMSD}}$ | top 1 $\mathrm{lDDT_{RNA}}$ | top 1 RMSD(Å) |
| RNArank | **0.770** | **0.823** | **0.675** | **0.873** | **0.577** | **0.644** | **0.435** | **73.1** | **9.6** |
| lociPARSE | 0.557 | 0.581 | 0.414 | 0.741 | 0.380 | 0.499 | 0.138 | 72.0 | 10.3 |
| RNA3DCNN | 0.130 | 0.108 | 0.142 | 0.587 | 0.099 | 0.256 | 0.049 | 56.9 | 15.6 |
| DFIRE-RNA | 0.128 | - | 0.459 | 0.546 | 0.410 | - | 0.199 | 68.5 | 12.9 |
| cgRNASP | 0.018 | - | 0.324 | 0.605 | 0.533 | - | 0.237 | 69.7 | 11.8 |
| cgRNASP-PC | 0.063 | - | 0.400 | 0.575 | 0.525 | - | 0.255 | 69.8 | 11.7 |
| cgRNASP-C | 0.094 | - | 0.423 | 0.570 | 0.551 | - | 0.272 | 66.7 | 13.2 |
| rsRNASP | 0.068 | - | 0.250 | 0.610 | 0.445 | - | 0.233 | 70.0 | 13.3 |
| Ideal value | 1 | 1 | 1 | 1 | 1 | 1 | 1 | 76.1 | 7.7 |

### Table S4. Performance on 12 CASP15 targets.

Values in **bold** indicate the best performance, while those with underlines represent the second-best. AUC was calculated on the overall data with an $\mathrm{lDDT_{RNA}}$ cutoff of 75 to differentiate between 'good' and 'bad' decoys. $\rho_{\mathrm{lDDT(Global)}}$, $\rho_{\mathrm{lDDT(Local)}}$ and $\rho_{\mathrm{RMSD}}$ represent the Spearman correlation between the assessment scores and the true global $\mathrm{lDDT_{RNA}}$, local $\mathrm{lDDT_{RNA}}$, and RMSD, respectively. All correlation values are reported as absolute values.

| **Method** | **Overall** | | | | **Per-target average** | | | | |
|---|---|---|---|---|---|---|---|---|---|
| | $\rho_{\mathrm{lDDT(Global)}}$ | $\rho_{\mathrm{lDDT(Local)}}$ | $\rho_{\mathrm{RMSD}}$ | AUC | $\rho_{\mathrm{lDDT(Global)}}$ | $\rho_{\mathrm{lDDT(Local)}}$ | $\rho_{\mathrm{RMSD}}$ | top 1 $\mathrm{lDDT_{RNA}}$ | top 1 RMSD(Å) |
| RNArank | **0.706** | **0.589** | **0.686** | **0.915** | 0.552 | **0.536** | **0.395** | **67.0** | **14.3** |
| lociPARSE | 0.511 | 0.505 | 0.200 | 0.673 | **0.574** | 0.419 | 0.304 | 64.6 | 19.3 |
| RNA3DCNN | 0.262 | 0.272 | 0.067 | 0.685 | 0.239 | 0.244 | 0.099 | 37.9 | 32.3 |
| DFIRE-RNA | 0.023 | - | 0.534 | 0.379 | 0.526 | - | 0.352 | 58.6 | 20.7 |
| cgRNASP | 0.096 | - | 0.511 | 0.404 | 0.475 | - | 0.290 | 63.8 | 19.9 |
| cgRNASP-PC | 0.092 | - | 0.525 | 0.384 | 0.414 | - | 0.266 | 56.1 | 25.7 |
| cgRNASP-C | 0.096 | - | 0.524 | 0.393 | 0.438 | - | 0.279 | 56.9 | 25.7 |
| rsRNASP | 0.023 | - | 0.434 | 0.446 | 0.573 | - | 0.347 | 64.0 | 19.6 |
| Ideal value | 1 | 1 | 1 | 1 | 1 | 1 | 1 | 73.8 | 7.7 |

### Table S5. Performance on 42 CASP16 targets.

Values in **bold** indicate the best performance, while those with underlines represent the second-best. AUC was calculated on the overall data with an $\mathrm{lDDT_{RNA}}$ cutoff of 75 to differentiate between 'good' and 'bad' decoys. $\rho_{\mathrm{lDDT(Global)}}$, $\rho_{\mathrm{lDDT(Local)}}$ and $\rho_{\mathrm{RMSD}}$ represent the Spearman correlation between the assessment scores and the true global $\mathrm{lDDT_{RNA}}$, local $\mathrm{lDDT_{RNA}}$, and RMSD, respectively. All correlation values are reported as absolute values.

| Method | Overall | | | | Per-target average | | | | |
|---|---|---|---|---|---|---|---|---|---|
| | $\rho_{\mathrm{lDDT(Global)}}$ | $\rho_{\mathrm{lDDT(Local)}}$ | $\rho_{\mathrm{RMSD}}$ | AUC | $\rho_{\mathrm{lDDT(Global)}}$ | $\rho_{\mathrm{lDDT(Local)}}$ | $\rho_{\mathrm{RMSD}}$ | top 1 $\mathrm{lDDT_{RNA}}$ | top 1 RMSD(Å) |
| RNArank | **0.778** | **0.507** | 0.722 | **0.934** | 0.549 | **0.421** | **0.385** | **60.0** | **25.8** |
| lociPARSE | 0.458 | <u>0.386</u> | 0.234 | <u>0.658</u> | **0.597** | <u>0.393</u> | <u>0.283</u> | <u>58.9</u> | <u>26.2</u> |
| RNA3DCNN | 0.252 | 0.253 | 0.019 | 0.584 | 0.478 | 0.298 | 0.175 | 51.7 | 32.0 |
| DFIRE-RNA | <u>0.503</u> | - | **0.751** | 0.230 | 0.531 | - | 0.201 | 57.5 | 29.6 |
| cgRNASP | 0.465 | - | 0.726 | 0.260 | 0.511 | - | 0.204 | 56.1 | 29.6 |
| cgRNASP-PC | 0.479 | - | 0.739 | 0.241 | 0.483 | - | 0.179 | 56.4 | 29.6 |
| cgRNASP-C | 0.490 | - | <u>0.751</u> | 0.241 | 0.485 | - | 0.161 | 51.8 | 31.9 |
| rsRNASP | 0.427 | - | 0.699 | 0.274 | <u>0.549</u> | - | 0.214 | 54.1 | 29.5 |
| Ideal value | 1 | 1 | 1 | 1 | 1 | 1 | 1 | 66.2 | 18.2 |

#### Table S6. Results for ablation study on T24 Targets.

Values in bold indicate the best performance, while those with underlines represent the second-best. AUC was calculated on the overall data with an $lDDT_{RNA}$ cutoff of 75 to differentiate between 'good' and 'bad' decoys. $\rho_{lDDT(Global)}$, $\rho_{lDDT(Local)}$ and $\rho_{RMSD}$ represent the Spearman correlation between the assessment scores and the true global $lDDT_{RNA}$, local $lDDT_{RNA}$, and RMSD, respectively. All correlation values are reported as absolute values.

| **Method** | **Overall** | | | | **Per-target average** | | | | |
|---|---|---|---|---|---|---|---|---|---|
| | $\rho_{lDDT(Global)}$ | $\rho_{lDDT(Local)}$ | $\rho_{RMSD}$ | AUC | $\rho_{lDDT(Global)}$ | $\rho_{lDDT(Local)}$ | $\rho_{RMSD}$ | top 1 $lDDT_{RNA}$ | top 1 RMSD(Å) |
| RNArank | <u>0.770</u> | **0.823** | 0.675 | 0.873 | **0.577** | **0.644** | <u>0.435</u> | **73.1** | **9.6** |
| w/o distance | 0.764 | 0.794 | 0.676 | 0.867 | 0.543 | 0.605 | 0.423 | <u>72.0</u> | 10.4 |
| w/o USR | **0.797** | 0.814 | <u>0.711</u> | **0.903** | 0.553 | 0.631 | **0.443** | 71.3 | 10.3 |
| w/o energy | 0.764 | 0.805 | 0.657 | <u>0.875</u> | <u>0.560</u> | <u>0.640</u> | 0.426 | 71.6 | <u>10.0</u> |
| w/o voxelization | 0.735 | 0.763 | **0.711** | 0.853 | 0.465 | 0.580 | 0.350 | 70.5 | 11.1 |
| Ideal value | 1 | 1 | 1 | 1 | 1 | 1 | 1 | 76.1 | 7.7 |

**Figure S1. Relationship between the two lDDT metrics and the other three established metrics for CASP15 targets.**

Each color corresponds to a target.

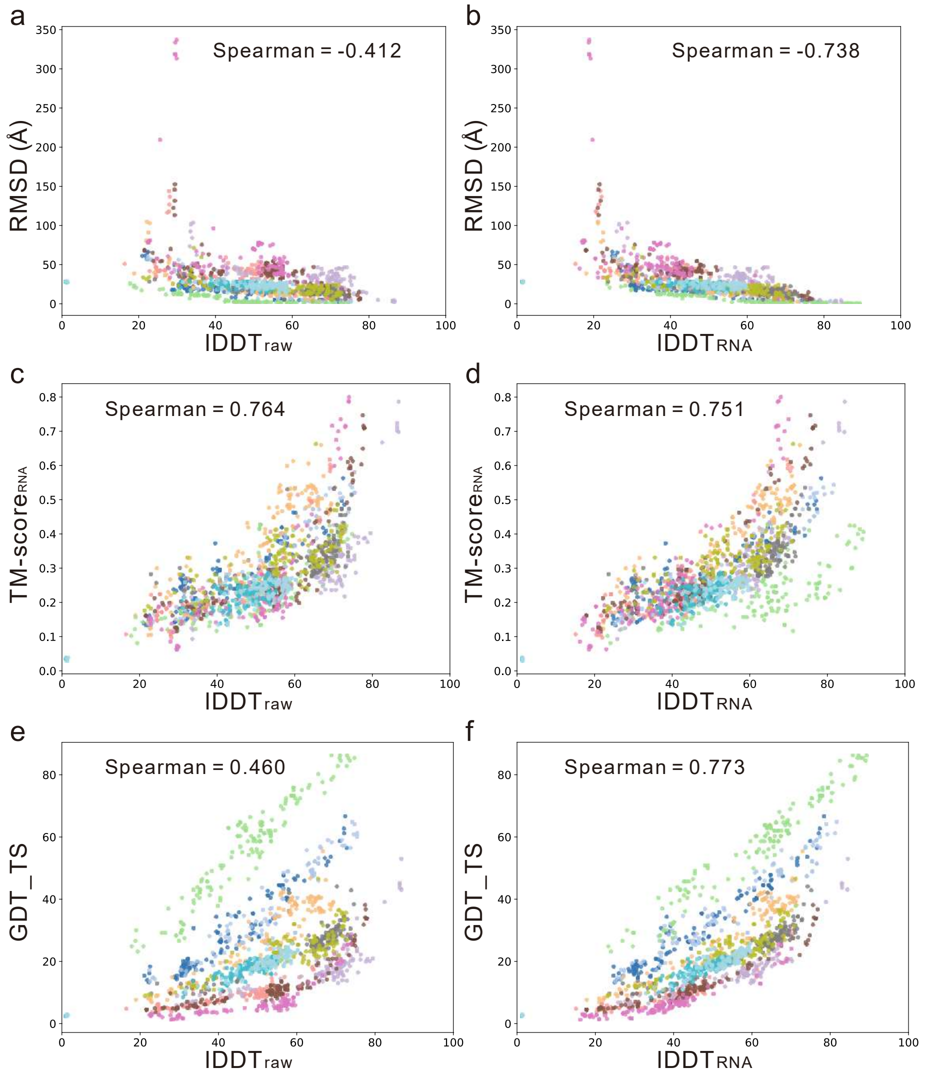

**Figure S2. Relationship between the two lDDT metrics and the other three established metrics for CASP16 targets.**

Each color corresponds to a target.

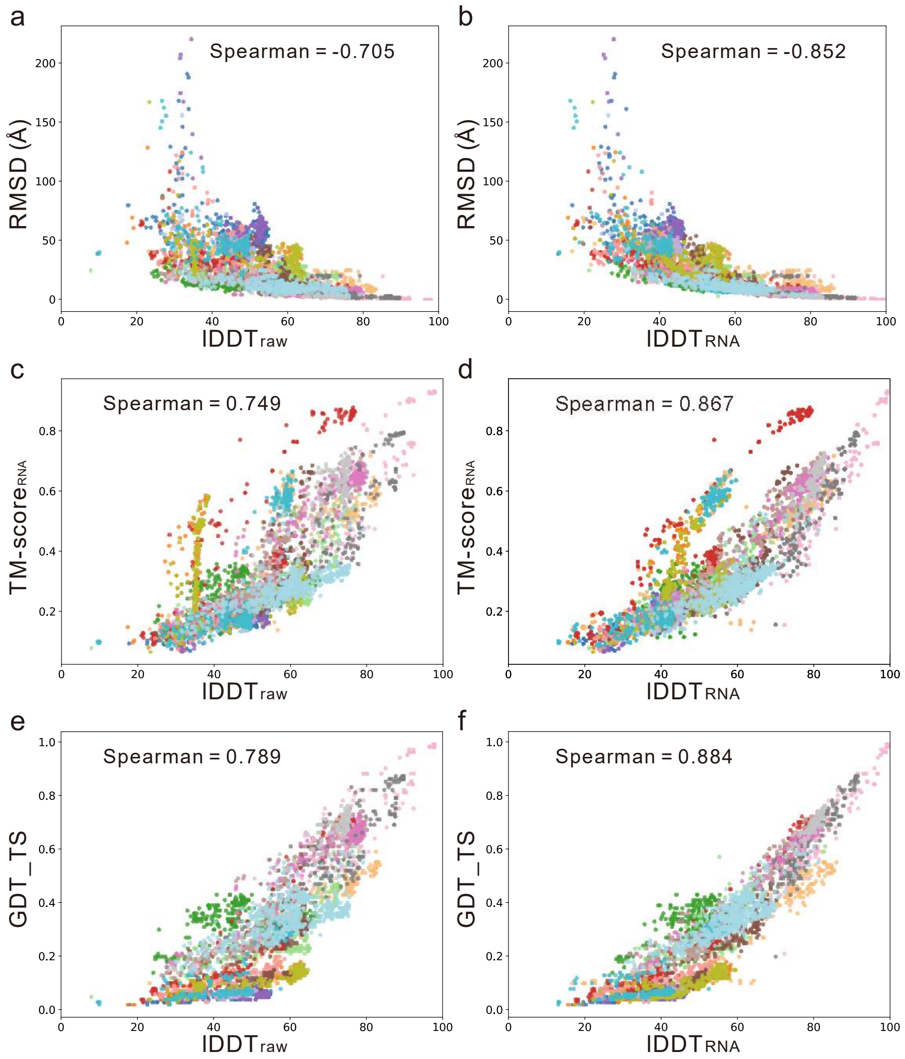

**Figure S3. Real RMSD-lDDT$_{RNA}$ relationships on three benchmark sets.**

**(a)** T24 dataset. **(b)** CASP15 dataset. **(c)** CASP16 dataset.

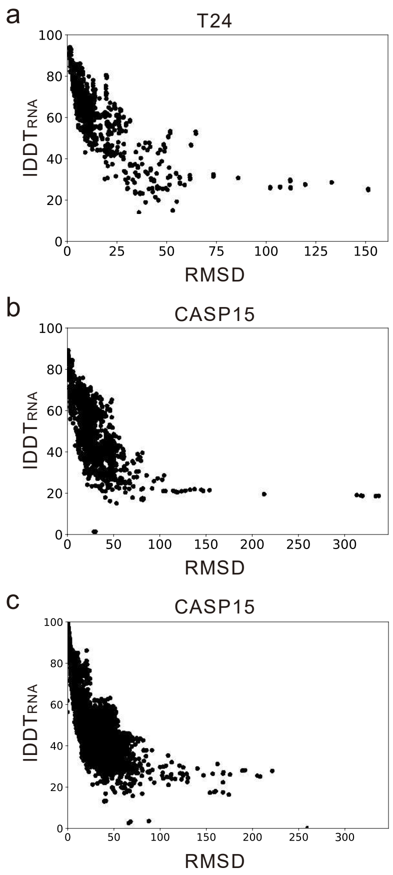

**Figure S4. ROC curves on two CASP datasets.**

**(a)** CASP15 dataset. **(b)** CASP16 dataset.

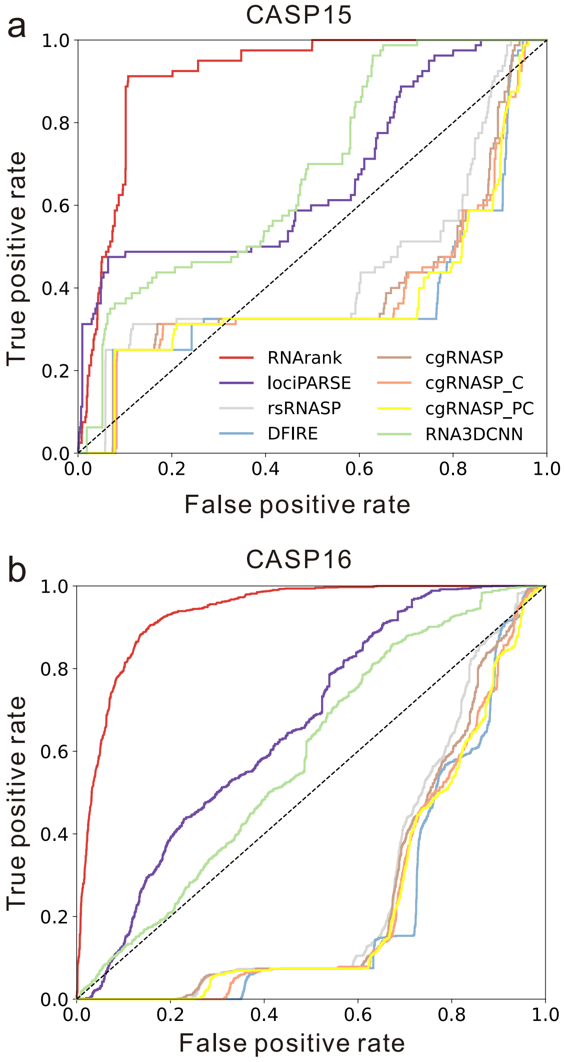

#### Figure S5. Real lDDT$_{RNA}$ vs. predicted scores on four CASP15 targets.

The red stars highlight the TS081 models.

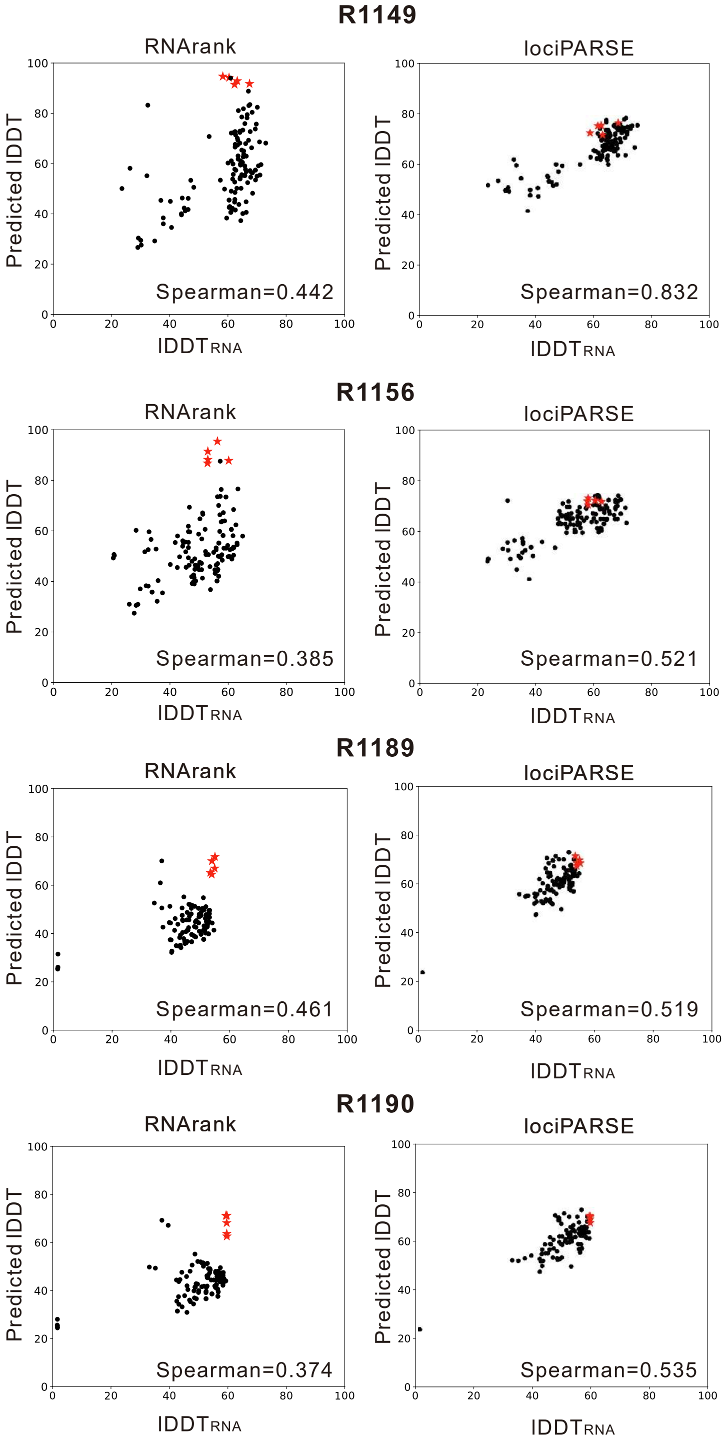

#### Figure S6. Best lDDT$_{RNA}$ score within the top-*k* ranked decoys on CASP16 dataset.

The vertical line indicates *k*=5.

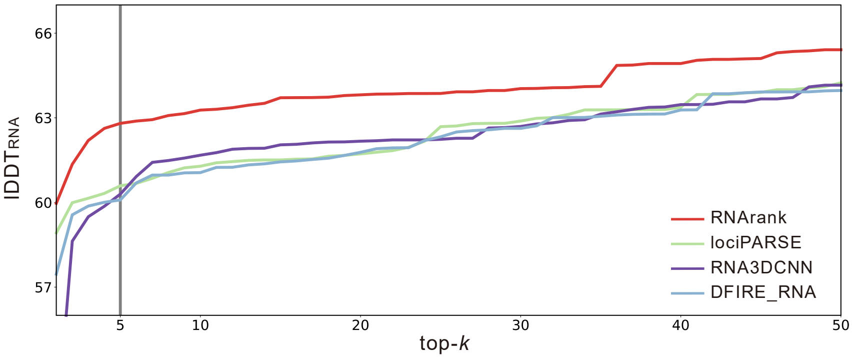

#### Figure S7. Length-dependency analysis of different scores on CASP16 dataset.

**(a)** The relationship between the ground-truth RMSD of decoy models and target length. **(b-g)** The relationship between the quality scores predicted by various methods and the target length. To minimize the impact of outliers, decoys with a $lDDT_{RNA}$ Z-score less than 0 (i.e., below average) were excluded from all panels.

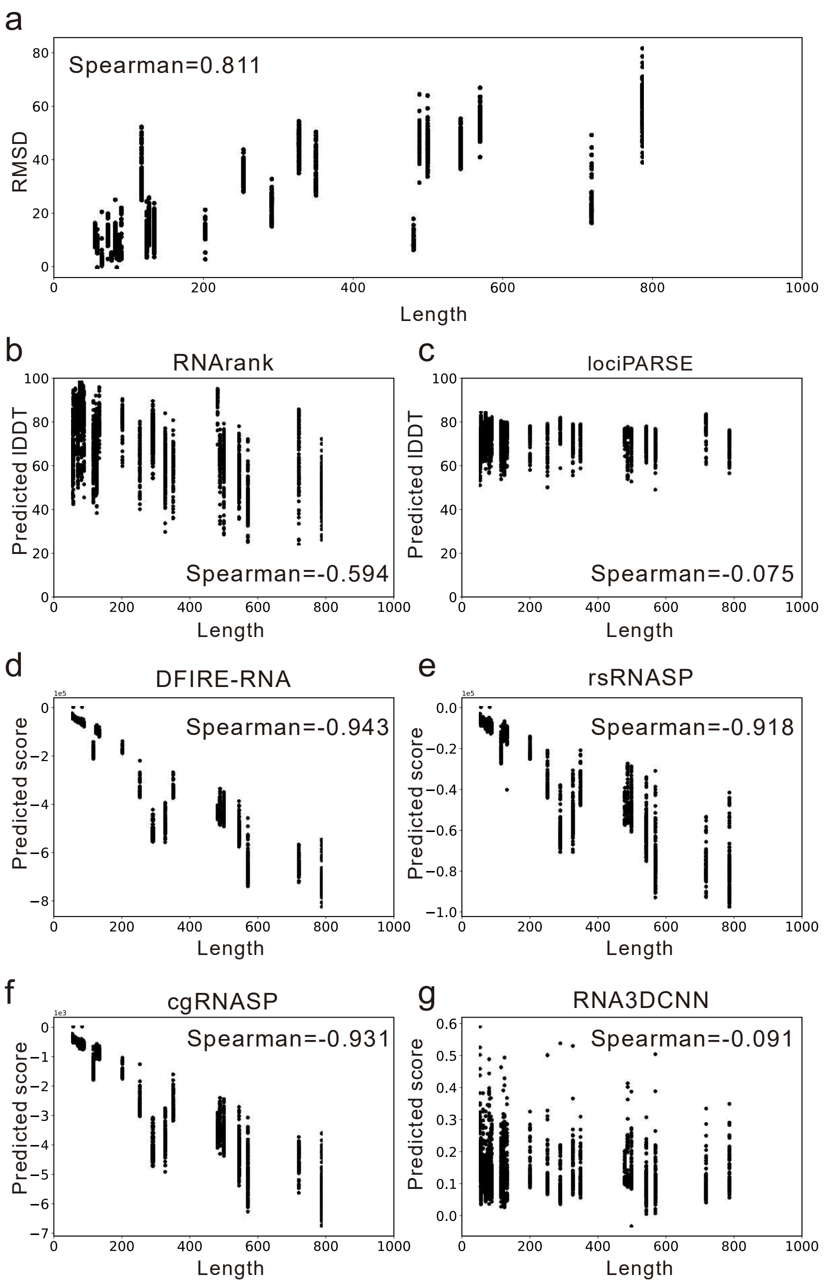

#### Supplementary References

1. Ballester PJ, Finn PW, Richards WG. Ultrafast shape recognition: Evaluating a new ligand-based virtual screening technology. *Journal of Molecular Graphics and Modelling* **27**, 836-845 (2009).

2. Guo SS, Liu J, Zhou XG, Zhang GJ. DeepUMQA: ultrafast shape recognition-based protein model quality assessment using deep learning. *Bioinformatics* **38**, 1895-1903 (2022).

3. Li J*, et al.* RNA3DCNN: Local and global quality assessments of RNA 3D structures using 3D deep convolutional neural networks. *PLOS Computational Biology* **14**, e1006514 (2018).

4. Abramson J*, et al.* Accurate structure prediction of biomolecular interactions with AlphaFold 3. *Nature* **630**, 493-500 (2024).

5. Pearce R, Omenn GS, Zhang Y. De Novo RNA Tertiary Structure Prediction at Atomic Resolution Using Geometric Potentials from Deep Learning. *bioRxiv*, 2022.2005.2015.491755 (2022).

6. Li Y, Zhang C, Feng C, Pearce R, Lydia Freddolino P, Zhang Y. Integrating end-to-end learning with deep geometrical potentials for ab initio RNA structure prediction. *Nat Commun* **14**, 5745 (2023).

7. Shen T*, et al.* Accurate RNA 3D structure prediction using a language model-based deep learning approach. *Nat Methods* **21**, 2287-2298 (2024).

8. Baek M, McHugh R, Anishchenko I, Jiang H, Baker D, DiMaio F. Accurate prediction of protein-nucleic acid complexes using RoseTTAFoldNA. *Nat Methods* **21**, 117-121 (2024).

9. Wang W*, et al.* trRosettaRNA: automated prediction of RNA 3D structure with transformer network. *Nat Commun* **14**, 7266 (2023).

10. Boniecki MJ*, et al.* SimRNA: a coarse-grained method for RNA folding simulations and 3D structure prediction. *Nucleic Acids Res* **44**, e63 (2016).
